# Integrated Proteogenomic Deep Sequencing and Analytics Accurately Identify Non-Canonical Peptides in Tumor Immunopeptidomes

**DOI:** 10.1101/758680

**Authors:** Chloe Chong, Markus Müller, HuiSong Pak, Dermot Harnett, Florian Huber, Delphine Grun, Marion Leleu, Aymeric Auger, Marion Arnaud, Brian J. Stevenson, Justine Michaux, Ilija Bilic, Antje Hirsekorn, Lorenzo Calviello, Laia Simó-Riudalbas, Evarist Planet, Jan Lubiński, Marta Bryśkiewicz, Maciej Wiznerowicz, Ioannis Xenarios, Lin Zhang, Didier Trono, Alexandre Harari, Uwe Ohler, George Coukos, Michal Bassani-Sternberg

**Affiliations:** Ludwig Cancer Research Center, University of Lausanne, Agora Center Bugnon 25A, 1005 Lausanne, Switzerland; Department of Oncology, Centre hospitalier universitaire vaudois (CHUV), Rue du Bugnon 46, 1005 Lausanne, Switzerland; Vital IT, Swiss Institute of Bioinformatics, 1015 Lausanne, Switzerland; Max Delbruck Centre for Molecular Medicine in the Helmholtz Association, Institute for Medical Systems Biology, Hannoversche Str 28, 10115 Berlin, Germany; École Polytechnique Fédérale de Lausanne (EPFL), Route Cantonale, 1015 Lausanne, Switzerland; School of Life Sciences, École Polytechnique Fédérale de Lausanne (EPFL), 1015 Lausanne, Switzerland; Swiss Institute of Bioinformatics, 1015 Lausanne, Switzerland; Department of Genetics and Pathology, International Hereditary Cancer Center, Pomeranian Medical University, Szczecin, Poland; International Institute for Molecular Oncology, 60-203 Poznań, Poland; University of Medical Sciences, 61-701 Poznań, Poland; Genome Center Health2030, Chemin de Mines 9, 1202 Genève, Switzerland; Department of Training and Research, CHUV/UNIL Agora Center Bugnon 25A, 1005 Lausanne, Switzerland; Center for Research on Reproduction and Women’s Health, University of Pennsylvania, Philadelphia, Pennsylvania 19104, USA; Department of Obstetrics and Gynecology, University of Pennsylvania, Philadelphia, Pennsylvania 19104, USA; Humboldt-Universitat zu Berlin, Departments of Biology and Computer Science, Unter den Linden 6, 10099 Berlin, Germany

## Abstract

Efforts to precisely identify tumor human leukocyte antigen (HLA) bound peptides capable of mediating T cell-based tumor rejection still face important challenges. Recent studies suggest that non-canonical tumor-specific HLA peptides that derive from annotated non-coding regions could elicit anti-tumor immune responses. However, sensitive and accurate mass-spectrometry (MS)-based proteogenomics approaches are required to robustly identify these non-canonical peptides. We present an MS-based analytical approach that characterizes the non-canonical tumor HLA peptide repertoire, by incorporating whole exome sequencing, bulk and single cell transcriptomics, ribosome profiling, and a combination of two MS/MS search tools. This approach results in the accurate identification of hundreds of shared and tumor-specific non-canonical HLA peptides and of an immunogenic peptide from a downstream reading frame in the melanoma stem cell marker gene ABCB5. It holds great promise for the discovery of novel cancer antigens for cancer immunotherapy.

## Introduction

The efficacy of T cell-based cancer immunotherapy relies on the recognition of HLA-bound peptides (HLAp) presented on the surface of cancer cells. Characterizing and classifying immunogenic epitopes is an ongoing endeavor for developing cancer vaccines and adoptive T cell-based immunotherapies. Neoantigens, which are peptides derived from mutated proteins, are absolutely tumor-specific yet mostly patient-specific and are implicated in the therapeutic efficacy of checkpoint blockade immunotherapy^1–4^. In contrast to tumor-specific neoantigens, tumor-associated antigens that are shared across patients may be more attractive for immunotherapy due to the more efficient and rapid treatment of a greater number of patients ^5–7^. Recent studies have focused on the discovery of non-canonical antigens, which are antigens derived from the translation of presumed non-coding transcripts. Such aberrant translation leads to the generation of peptide sequences that are missing in conventional protein sequence repositories and are therefore considered novel^8, 9^. If such translation events lead to the presentation of novel and immunogenic HLA ligands, then these occurrences could substantially expand the repertoire of targetable epitopes for cancer immunotherapy^9–20^. Currently, approximately 1% of the entire genome is annotated as protein-coding regions, yet, 75% of the genome can be transcribed and theoretically translated, potentially offering a pool of novel peptide targets^21^.

To date, the only analytical methodology allowing the direct identification of the *in vivo* presented HLAp repertoire is mass spectrometry (MS)^22^. Often, MS-based immunopeptidomic discoveries are limited to the standard, available protein sequence database, usually containing only annotated protein-coding sequences. Recently, several studies have included protein sequences derived from the translation of transcripts identified from RNA-Seq, or from ribosome profiling, into MS-based searches^9, 23–28^. Overall, these studies warrant further development in many key aspects: Importantly, elevated false discovery rates (FDRs) for the non-canonical space can occur when MS reference data are populated with protein sequences derived from all potential three- or six-frame translations of transcribed regions^29^. Several studies did not compute FDRs or applied sample-specific thresholds for FDR calculations^24, 28^. Furthermore, rigorous experimental confirmation with targeted MS for such non-canonical sequences is currently lacking. Also, current workflows often introduce a risk of bias by pre-filtering peptide identifications based on HLA-binding predictions^24, 28^. Finally, the overall biogenesis of non-canonical HLA binding peptides (noncHLAp) remains understudied due to *a-priori* restriction of the search space to tumor-specific non-canonical protein sequences^24^.

Here we describe a proteogenomic approach to identify tumor-specific noncHLAp derived from translation of presumed non-coding transcripts, such as from (long) non-coding genes (lncRNAs), pseudogenes, untranslated regions (UTRs) of coding genes, and transposable elements (TEs). We performed immunopeptidomics and integrated in our analyses tumor exome, bulk and single cell transcriptome, and whole translatome data. We then implemented NewAnce, a <u>new an</u>alytical approach for <u>n</u>on-<u>c</u>anonical <u>e</u>lement identification that combines two MS/MS search tools, along with group-specific FDR calculations to identify noncHLAp. Together, this unveiled an unprecedentedly large number of novel noncHLAp, highlighting the potential of this approach to enlarge the range of targetable epitopes in cancer immunotherapy.

## Results

### A comprehensive strategy for noncHLAp identification

MS-based immunopeptidomics was performed on seven patient-derived melanoma cell lines and two pairs of lung cancer samples with matched normal tissues (Fig. 1a). This resulted in the identification of 60,320 unique protein-coding HLA class I bound peptides (protHLAIp) and 11,256 protein-coding HLA class II-bound peptides (protHLAIIp). For the exploration and validation of *in vivo* naturally presented non-canonical peptides, whole exome and RNA-Seq data were generated for all samples (Fig. 1a **and Supplementary Table 1**). For every sample, we extracted from RNA-Seq data expressed non-canonical genes such as lncRNAs and pseudogenes. In addition, we applied an analytical pipeline to assign TE-derived RNA-Seq reads to single loci (See Methods section for more details), resulting in expression data for transcribed TEs. All the above transcripts were subsequently translated in three forward open reading frames (ORFs) (stop-to-stop). For every sample, the novel *in-silico* translated protein sequences were concatenated to a personalized canonical proteome reference containing allelic variant information from patient tumor exome data. Finally, we searched the MS immunopeptidomics data against these personalized reference databases.

**Fig. 1.**
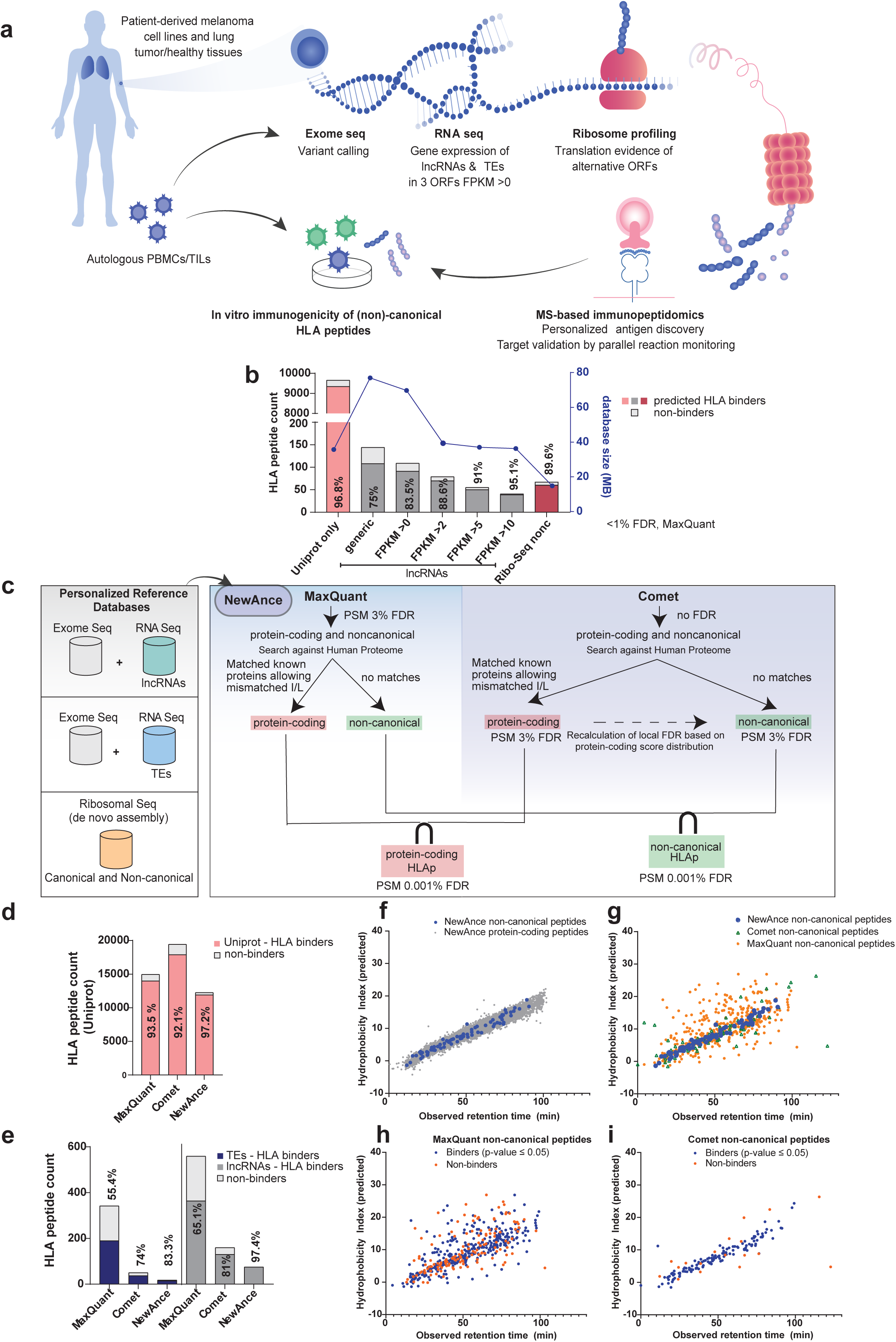
A novel proteogenomics approach for the robust identification of noncHLAp. **a** A schematic of the entire workflow is shown, where tumor tissue samples or tumor cell lines were obtained from patients, and exome, RNA and Ribo-Seq performed to provide a framework to interrogate the non-canonical antigen repertoire. HLAp are immunoaffinity purified from cancer cell lines and matched tumor/healthy lung tissues and analyzed by MS. Immunopeptidomics spectra were then searched against RNA and Ribo-Seq based personalized protein sequence databases that contain non-canonical protein sequences. MS-identified noncHLAIp were validated by targeted MS-based PRM and tested for immunogenicity using autologous T cells or PBMCs. **b** The percentage of HLA-binders with a MixMHCpred p-value < 0.05 is used to evaluate the accuracy of the identified HLAIp as a function of database size (blue line). Percentage of binders obtained for each condition is shown for each bar, for melanoma cell line 0D5P. **c** Different protein sequence databases combining whole exome sequencing, and inferred from RNA-Seq and Ribo-Seq data were utilized. NewAnce was implemented by retaining the PSM intersection of the two MS search tools MaxQuant and Comet, and applying group-specific FDR calculations for protHLAp and noncHLAp. **d** The percentages of protein-coding HLA-I binders were assessed for 0D5P for each MS search tool (MaxQuant and Comet at FDR 3%) and NewAnce. **e** Similar to d, the comparisons were performed for the different non-canonical antigen classes. **f** Retention predictions for peptides identified with melanoma 0D5P. Observed mean retention time is plotted against hydrophobicity indices for NewAnce identified protein coding versus non-canonical peptides. **g** All peptides identified with each tool (MaxQuant, Comet, NewAnce) were analyzed based on their hydrophobicity indices. **h** MaxQuant or **i** Comet identified 8-14 mer peptides were analyzed based on their HLA binding predictions which were assessed with MixMHCpred.

### Database size affects the level of false positives in noncHLAp identifications

In silico translation of transcripts in three reading frames results in a large number of potential protein sequences. In proteogenomics, searching MS data against such inflated protein reference databases may propagate false positives ^29, 30^. Hence, our first investigative step was to understand the impact of database size on the level of false positives in immunopeptidomics datasets. We searched reference databases containing canonical (i.e. UniProt) and our non-canonical protein sequences with a single search tool (MaxQuant) and a global 1% FDR. The accuracy was assessed by assigning HLA-binding prediction scores to the MS-identified HLAIp with MixMHCpred^31^. We reasoned that non-canonical HLA class I bound peptides (noncHLAIp) should follow the same binding rules as protHLAIp^32^. First, we compared a generic non-canonical protein sequence database derived from the three forward frame (“three-frame”) translation of all annotated non-coding genes from GENCODE^33^ with a sample-specific protein sequence database derived from the three-frame translation of lncRNAs and pseudogenes from the RNA-Seq data, using an expression cut –off value of FPKM (fragments per kilobase of transcript per million mapped reads >0). Additional databases of decreasing size were assembled, by retaining only those sequences that originated from more highly expressed genes (FPKM >2, >5 or >10). Reducing the size of the database by personalizing and focusing on highly expressed genes led to an increase in the percentage of noncHLAIp that were predicted to bind to their respective HLA alleles (MixMHCpred p-value<0.05) (Fig. 1b).

### NewAnce improves accurate identification of hundreds of noncHLAp

Restricting the database to protein sequences originating from highly expressed genes should on one hand improve the accuracy of MS based non-canonical peptide identification, while on the other hand lead to the potential loss of peptides coming from lower expressed transcripts. To circumvent the need to exclude protein sequences based on low expressed transcripts, we developed the computational module called NewAnce, which combines the MS search tools MaxQuant^34^ and Comet^35^, with the implementation of a group-specific strategy for FDR calculation. All HLAp identified by either of the search tools were consequently matched against an up-to-date UniProt sequence database (95,106 protein sequences, with isoforms) to extract novel noncHLAp that do not map back to known human proteins in UniProt. For every sample, FDRs were calculated separately for the protHLAp and noncHLAp (Fig. 1c **and Supplementary Fig. 1a**). Only consensus (intersection) peptide-spectrum-matches (PSMs) from Comet and MaxQuant were retained for further downstream analyses. As most false positive PSMs are specific to one search tool, NewAnce led to an estimated FDR of <0.001%.

With NewAnce, the number of protHLAIp identified across 11 samples ranged from 3,490 to 16,672 per sample, and from 817 to 5,777 for protHLAIIp (**Supplementary Table 2**). Furthermore, up to 148 noncHLAIp per individual sample were identified with NewAnce, with a combined total of 452 unique noncHLAIp (**Supplementary Table 2 and Supplementary Data 1**). Over the four HLA-II expressing samples investigated, only 4 lncRNA derived noncHLAIIp out of 11,256 protHLAIIp were detected.

We employed two complementary methods to assess the accuracy of our approach. First, we predicted the binding of peptides to their respective HLA allotypes. Across all 11 samples, 90% of the noncHLAIp and 91% of the protHLAIp identified with NewAnce were predicted to bind the HLA allotypes (median values, **Supplementary Fig. 1b**). As expected, NewAnce detected less HLAp than Comet (PSM FDR of 3%) or MaxQuant (PSM FDR of 3%) and more HLAp when the routinely applied FDR of 1% was applied by MaxQuant alone (Fig. 1d, **Supplementary Fig. 1b-f and Supplementary Table 2**). Importantly, for the noncHLAIp repertoire (lncRNAs, pseudogenes and TEs), significantly higher percentages of peptides predicted to bind the HLA allotypes were identified by NewAnce compared with those identified by MaxQuant or Comet alone (Fig. 1e **and Supplementary Fig. 1b-f**).

In addition, we correlated the observed mean retention time (RT) of a given peptide against the predicted hydrophobicity index (HI), which corresponds to the percentage of acetonitrile at which the peptide elutes from the analytical HPLC system. Predicting the sequence specific hydrophobicity indices of peptides identified by NewAnce showed that the RT distribution of non-canonical peptides was on the diagonal line, and was not significantly different than the distribution of protein-coding peptides, supporting their correct identification (Fig. 1f) (one sided F-test p-value:1.0e+0). However, we observed a significant difference in RT distribution when comparing non-canonical peptides from NewAnce to those identified by MaxQuant (one sided F-test p-value: 6.3e−32) or Comet alone (one sided F-test p-value: 8.4e−20) (Fig. 1g).

Moreover, a commonly applied approach to boost non-canonical peptide identifications would be to search the MS data with a single tool (or a union of two tools) applying a permissive FDR followed by an additional step of filtering to include only peptides predicted to bind the relevant HLA allotypes^36^. To evaluate this approach, we compared the correlation between HI and RT of predicted non-canonical HLA binders and non-binders identified at 3% PSM FDR with either MaxQuant (Fig. 1h) or Comet (Fig. 1i). Predicted binders showed better correlations between HI and RT compared to non-binders (one sided F-test p-values 8.4e-6 for MaxQuant and 4.4e-18 for Comet). These correlations where fairly poor for MaxQuant, while a much better correlation was calculated for Comet likely due to the conservative group specific FDR control strategy we applied for Comet. In conclusion, these comprehensive analyses underline the superiority of NewAnce over the above alternatives.

Notably, when examining the source protein sequence origin of all noncHLAIp, we detected an enrichment towards the C-terminus of their precursor protein sequences. This effect was also observed for protHLAIp originating from similarly short canonical proteins (**Supplementary Fig. 2a-b**).

### MS-targeted validation and Ribo-Seq confirm a fraction of noncHLAIp

To experimentally validate the NewAnce computational pipeline, we investigated a selection of NewAnce-identified HLAp from a melanoma sample (0D5P) with targeted MS-based analyses. All identified noncHLAIp (lncRNAs and TEs, n=93), as well as a similar-sized subset of protHLAIp from clinically relevant tumor-associated antigens (TAAs, n=71) detected in 0D5P were synthesized in their heavy isotope-labelled forms for MS-targeted validation. The selected TAAs were chosen solely based on their interesting tumor-associated biological functions, as these are known cancer/testis or melanoma antigens. Here, MS-based targeted confirmation with parallel reaction monitoring (PRM) was directly compared between the non-canonical and canonical peptide groups by spiking the heavy-labelled peptides into multiple independent replicates of 0D5P immunopeptidomic samples. This revealed that confirmation is superior for protHLAIp than for noncHLAIp (78.5% for TAAs versus 55.2% for lncRNAs and 27.7% for TEs) (Fig. 2a, **Supplementary Table 3 and Supplementary Data 2**). We also observed that PRM validation is dependent on source RNA expression level (**Supplementary Fig. 3a-d**), measured peptide intensities (**Supplementary Fig. 3e-h**), and detectability by MS/MS across multiple 0D5P replicates (**Supplementary Fig. 3i-l**).

**Fig. 2.**
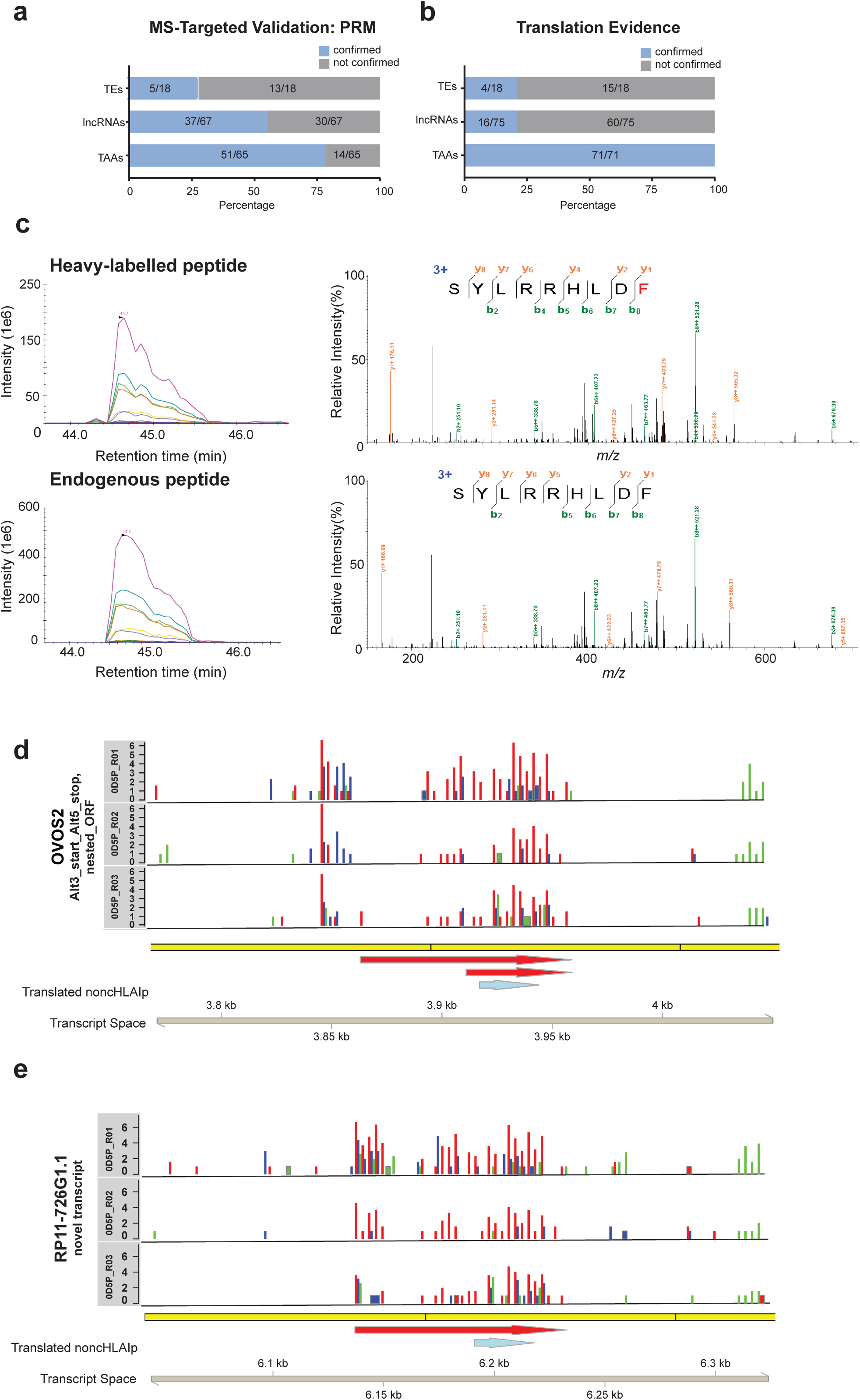
MS-based targeted validation by PRM and ribosome footprint pattern as evidence of non-canonical peptide generation. A set of protein-coding tumor-associated antigens and noncHLAIp (lncRNAs and TEs) from melanoma 0D5P were synthesized in its heavy labelled form and spiked back into replicates of eluted HLAIp from 0D5P to confirm the presence of the endogenous HLAIp. The proportions of confirmed and non-confirmed HLAIp by **a** PRM and **b** through Ribo-Seq targeted validation are shown for each of the antigen classes. **c** An example of co-elution profiles of transitions of heavy labelled and endogenous noncHLAIp (from lncRNA; SYLRRHLDF) from 0D5P (left) is shown. MS/MS fragmentation pattern further confirms the presence the endogenous peptide (Δm=10Da) (right). **d, e** The Ribo-Seq profiles for two source genes show the frequency of Ribo-Seq reads on ribosome’s P-site in three replicates. Library size-normalized P-sites per basepair are shown on a log2 scale on the Y-axis, with P-sites inferred as a constant offset from the 5’ end of the footprint, for each read length. Colored bars represent different reading frames. Yellow bars below the plots represent exons. For example, the noncHLAIp SYLRRHLDF in OVOS2 (blue arrow) falls within two nested, Ribo-Seq-supported ORFs (red arrows), within which most P-sites (red bars) fall in the first reading frame.

As a further targeted validation strategy for the noncHLAIp, Ribo-Seq, which involves the sequencing of ribosome protected fragments (RPFs), was performed on the sample 0D5P. Periodic RPF distributions (see Methods section) that supported translation in the correct ORF of transcript encoding the identified noncHLAIp was observed for 22.2% of TE peptides and 21.3% of lncRNA peptides, compared to 100% of the TAAs (Fig. 2b). Notably, nine lncRNA HLAIp, and two TE peptides were validated by both PRM and by Ribo-Seq approaches. For example, the noncHLAIp SYLRRHLDF was confirmed by MS (Fig. 2c), and the translated ORF that generated the peptide was mapped back to two novel transcripts (Fig. 2d-e).

### Low RNA expression level is a limiting step in noncHLAIp presentation

We then characterized in more depth the expression levels of source RNAs encoding the HLAIp. For this purpose, we compared all identified source genes of protHLAIp to source genes of noncHLAIp in the 0D5P sample. The protein-coding source genes had a median FPKM value of 9.3, whereas the non-canonical source genes showed overall lower expression, with a median FPKM of 2.1 (Fig. 3a-b). Generally, the number of unique peptides identified per gene increased with higher levels of expression. PRM-validated noncHLAIp covered a large and dynamic range of gene expression, and interestingly, a few were confirmed at very low source RNA expression levels (Fig. 3c-d).

**Fig. 3.**
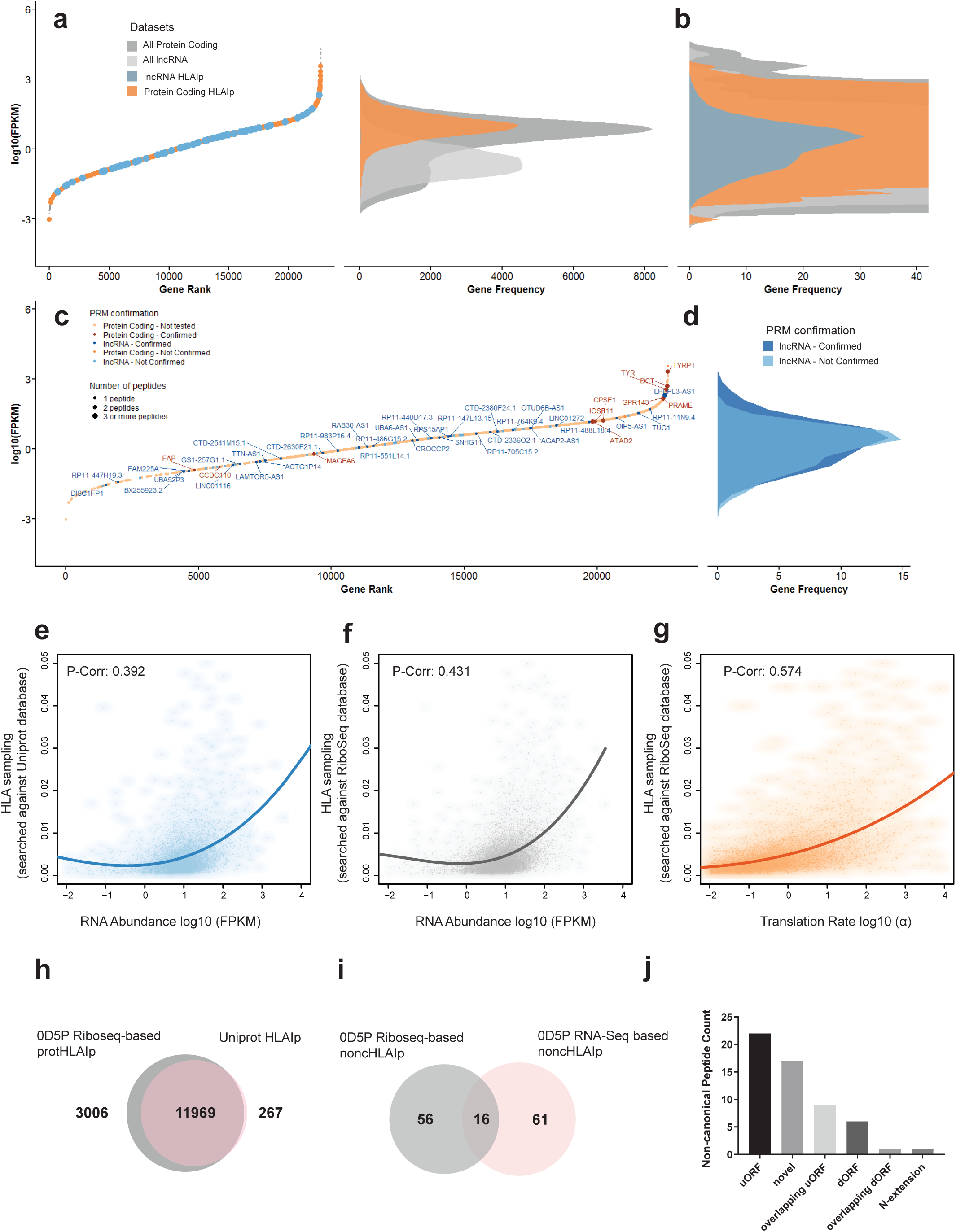
RNA-Seq and Ribo-Seq based gene expression analyses for the characterization of the protein-coding and non-canonical HLA immunopeptidome for melanoma 0D5P. **a** Genes are ranked based on RNA expression in 0D5P. P protein-coding (orange) and non-coding (blue) source genes, in which HLAIp were identified. The frequency distribution of gene expression for protein-coding and non-coding (lncRNA) genes is shown. **b** Magnification of the region of interest to show the distribution of noncHLAIp source gene expression. **c** Source gene restricted plot. Targeted MS validation was performed and its confirmation denoted for all identified non-canonical and a subset of protHLAIp (selected TAAs). Confirmed hits indicate that one or more peptides from that source gene were validated by PRM. Point sizes represent the number of peptides identified per source gene. **d** Frequency distribution of gene expression for MS confirmed versus non confirmed (or inconclusive) noncHLAIp. Scatterplots show the correlation between **e** UniProt based HLA-I sampling and RNA abundance, **f** Ribo-Seq-based HLA-I sampling and RNA abundance, and **g** Ribo-Seq-based HLA-I sampling and translation abundance. HLA-I sampling was calculated from adjusted peptide counts normalized by protein length. Determination of correlation between gene expression and HLA-I sampling was assessed by fitting a polynomial curve of degree 3 to each dataset. Pearson correlations were calculated to assess the correlation between the fitted curve and the corresponding dataset. **h** With data derived from 0D5P, a comparison of the overall overlap in unique HLAIp identifications with RNA-Seq based and Ribo-Seq based assembled databases for MS search is shown. **i** Overlap of noncHLAIp identifications found with RNA-Seq and Ribo-Seq based searches. **j** The number of identified noncHLAIp by Ribo-Seq is depicted with their respective source gene types.

The lower levels of expression of source genes that generated noncHLAp prompted us to investigate the regulation of non-canonical HLA presentation, and whether this can be induced with drug treatments. We treated melanoma cells either with decitabine, a DNA methyltransferase inhibitor, known to reactivate epigenetically silenced genes, or with IFN gamma (IFNγ), known to upregulate antigen presentation^37–40^. As expected, when T1185B melanoma cells were treated with IFNγ, we observed large quantitative changes in the presentation of protHLAIp. Specifically, enhanced presentation of peptides derived from immune-related genes was observed, likely due to their higher gene expression and the increase in the production of HLA-I molecules (**Supplementary Fig. 3m**). However, no obvious change was observed for the noncHLAIp repertoire, with 60% of the identified noncHLAIp remaining unaltered upon IFNγ treatment, suggesting that transcription is the limiting step in presentation of noncHLAIp (**Supplementary Fig. 3n**). Furthermore, we explored the effect of hypomethylating agent decitabine on noncHLAIp in melanoma. Although decitabine induced expression of selected hypomethylating-induced immune genes^41^, TAAs and non-canonical transcripts (**Supplementary Fig. 3o-q**), changes in the 0D5P noncHLAIp repertoire were modest (data not shown). Nonetheless, we identified and confirmed the presence of a unique decitabine -induced noncHLAIp derived from a lncRNA (**Supplementary Fig. 3r**).

### Integration of Ribo-Seq data improves the coverage of immunopeptidomes and identification of additional noncHLAIp

Next, we hypothesized that immunopeptidomes would be better associated with translatomes than transcriptomes. To build the translatome-based database for MS search, all ORFs showing periodic RPF distribution were extracted for the 0D5P sample and the transcribed frames were translated *in-silico*. This technique reduced the size of the search space, and we used this independent discovery method in our study in order to identify additional noncHLAIp, including those from novel ORF in coding genes.

We investigated to what extent a protein sequence database inferred by Ribo-Seq could replace the search performed with protein sequences derived from three-frame translation of expressed RNA species. Using 0D5P as a representative immunopeptidomic dataset, we observed a positive correlation between RNA expression and HLAIp sampling (see Methods section) searched against a canonical protein sequence database (r= 0.392) (Fig. 3e). Then we searched the same immunopeptidomics MS data against the *de novo* assembled Ribo-Seq inferred database, and we correlated this HLAIp sampling with RNA abundance (Fig. 3f) or with translation rates based on the spectral coefficient of 3-periodic signal in Ribo-Seq data (see Methods section) (Fig. 3g). This resulted in a significantly higher positive correlation between HLAIp sampling searched against a Ribo-Seq inferred database and translation rate (r=0.574) than with the overall RNA abundance (r=0.431, two-sided p-value<10e-16). Thus, there is evidence that the immunopeptidome, at least for 0D5P sample, is better captured by the translatome than the transcriptome.

Notably, restricting the database to actual translation products by Ribo-Seq provided a deeper coverage of the immunopeptidome than a canonical protein sequence database (Fig. 3h). This led to the identification of additional noncHLAIp derived from ORFs originating in 5’ or 3’ untranslated regions, presumed non-coding RNAs, retained introns, and pseudogenes, with the majority coming from either annotated upstream or entirely novel ORFs (Fig. 3i-j). Many of these additional noncHLAIp were missed using the RNA-Seq inferred database. Of note, this method also takes into account products arising from ribosomal frameshifting, which could be relevant in the context of non-canonical antigens^42^. Interestingly, only 16 common lncRNA-derived noncHLAIp were found when comparing both strategies. This likely reflects the limited detection of periodic Ribo-Seq reads in transcripts with low expression, or low mappability (**Supplementary Fig. 4a**).

### Single-cell transcriptomics reveals transcriptional heterogeneity of presumed non-coding genes

Tumor cell heterogeneity could present one of the key factors for immune escape leading to the inefficacy of cancer immunotherapies. In an attempt to understand the pattern of non-coding gene expression at the single cell level, we performed single-cell RNA-Seq (scRNA-Seq) on the 0D5P melanoma cell line. Overall, 1,400 cells were sequenced at a total depth of 176 million reads resulting in the detection of a median of 6,261 genes per cell (total of 19,178 detected genes). As expected, clustering of 0D5P cells revealed dependency on the cell cycle status (Fig. 4a) and source genes associated with cell cycle could be explored (Fig. 4b-c). First, we confirmed that the antigen presentation machinery was uniformly expressed in all cells, as well as many of the selected TAAs (Fig. 4d).

**Fig. 4.**
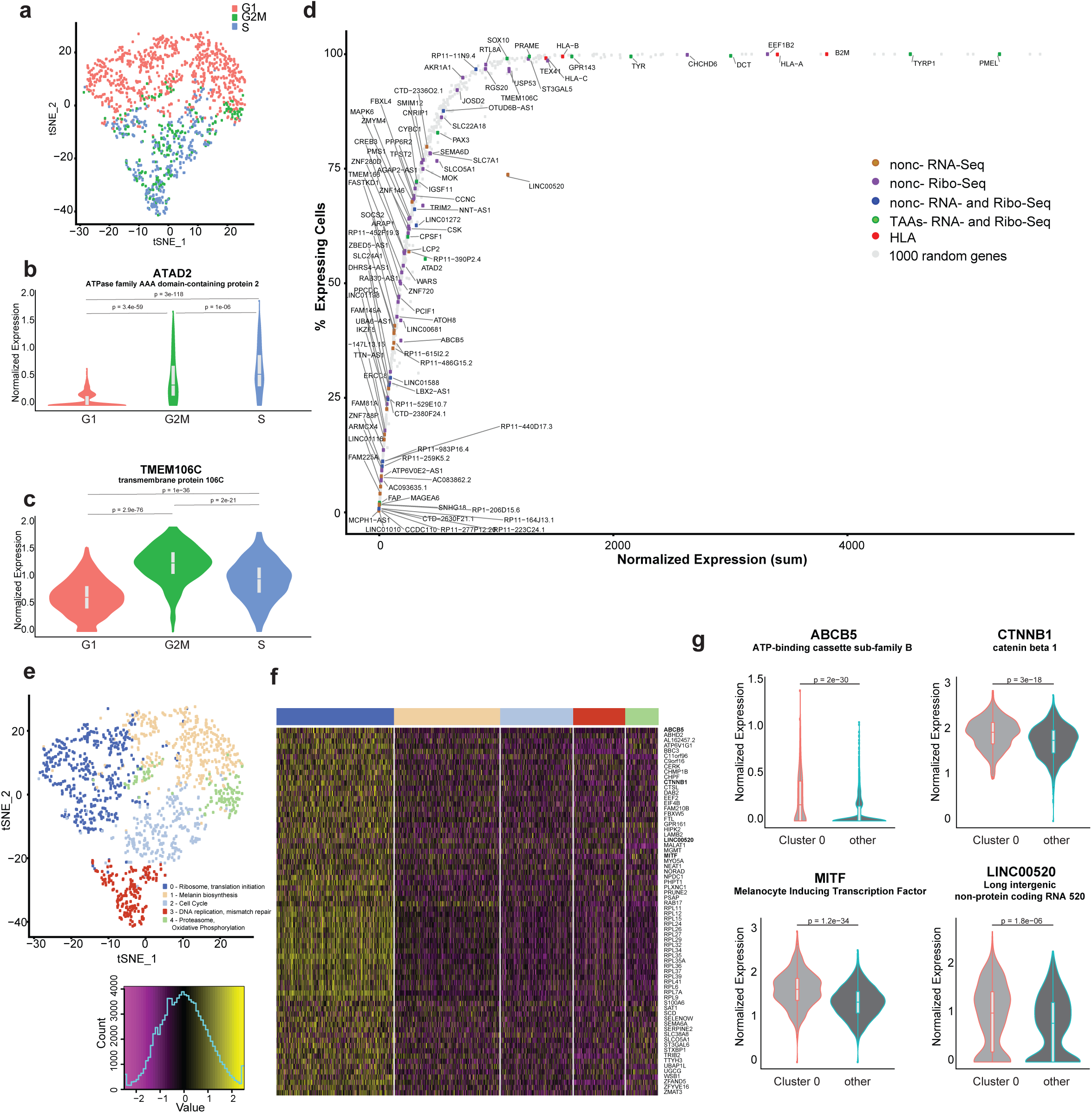
ScRNA-Seq reveals non-coding transcriptional heterogeneity in melanoma 0D5P. **a** t-SNE plot of the 1,400 cells colored by their “cell cycle” scores. **b** Examples of genes that are cell-cycle dependent: ATAD2, a tumor associated antigen, and **c** TMEM106C, where a noncHLAIp originated from. **d** Genes of interest were plotted based on their sum normalized expression by scRNA-Seq and ordered based on percentage of cells that expressed the gene. Color codes denote the type of HLAIp identified from those genes. For clarity purposes, only some genes were labelled. **e** t-SNE plot of the 1,365 cells colored by the five identified clusters. Clusters were annotated based on functional enrichment analyses of marker genes. **f** Heatmap showing the scaled and centered expressions of marker genes for cluster 0. Cluster colors from (e) are represented above the plot. **g** Expression profiles of four marker genes for cluster 0 over all other clusters, for two well-known cancer biomarkers MITF and CTNNB1, and two source genes where noncHLAIp were identified: ABCB5 gene with a dORF, and LINC00520. The p-values represented in (b), (c) and (g) were obtained with Wilcoxon tests.

Out of the 71 non-canonical source genes identified by bulk transcriptomics, 35 were detected also at the single-cell level (Fig. 4d). HLAIp derived from non-canonical source genes were detected with higher coverage at the single cell were those confirmed by PRM (6 out of 8 genes confirmed in >50% cells, and 14 out of 27 genes in <50% cells) and by Ribo-Seq (37 out of 41 genes confirmed in >50% cells, and 25 out of 46 genes in <50% cells) (Fig. 4d, **Supplementary Fig. 4b-c**). Profiles of source non-canonical genes clearly show expression heterogeneity and nearly none of them were uniformly expressed across cells, though the limited sensitivity of scRNA-Seq could account for this variation. Expression of LINC00520 was higher than expected given its detection in only 75% of cells, suggesting that it is not uniformly expressed (Fig. 4d).

We further explored marker genes associated with a cluster of 0D5P cells significantly expressing LINC00520 (Fig. 4e-f). Interestingly, we found that LINC00520 was co-expressed with the ATP-binding cassette sub-family B member 5 (ABCB5) gene, that mediates chemotherapeutic drug resistance in stem-like tumor cell subpopulations in human malignant melanoma and is commonly over-expressed on circulating melanoma tumor cells^43^, with beta-catenin (CTNNB1) which is a key regulator of melanoma cell growth^44^, and with its critical downstream target the microphthalmia-associated transcription factor (MITF) that mediates melanocytes differentiation^45^ (Fig. 4g). The source RNA expression of ABCB5 was detected in only 37% of 0D5P cells, however, as shown below, the ABCB5 gene encodes a novel ORF that gives rise to an immunogenic epitope. Indeed, immune-targeting of non-canonical targets expressed on a subset of aggressive or melanoma stem cells could be beneficial.

### Identification of tumor-specific noncHLAIp

As our initial MS search space was not restricted to protein sequences derived from tumor-specific transcripts, we investigated in retrospect the potential of identified noncHLAIp to be classified as tumor specific. A public database of RNA sequencing data from 30 different healthy tissues (GTEx^46^) was assessed at a strict 90^th^ percentile, which represents the expression value cut-off for the top 10% expressed genes. We identified 335 noncHLAIp from 280 lncRNA genes in the seven melanoma samples of which 23% were expressed only in our tumor samples, and not in any of the healthy tissues (excluding testis due to its immune-privileged nature) (Fig. 5a). Among these was the tumor-specific LINC00518 gene that has been proposed as a two-gene classifier for melanoma detection together with the tumor associated antigen PRAME^47^.

**Fig. 5.**
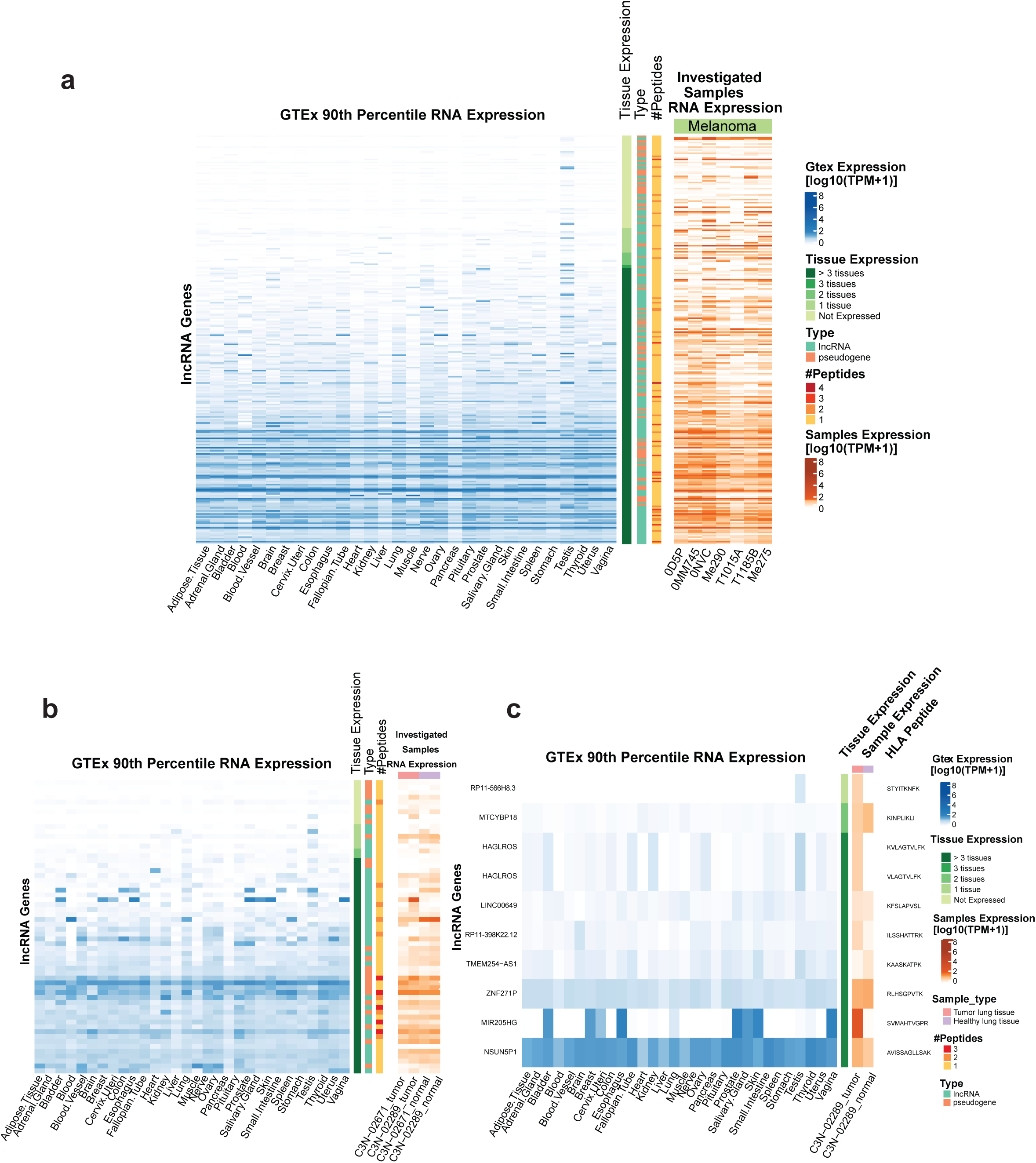
A comparison of non-coding source gene expression of investigated samples to healthy tissues (GTEx) reveals that a substantial proportion of source non-coding genes are tumor-specific. **a** The heatmap of lncRNA source genes shows the 90th percentile gene expression over 30 healthy tissues on the left, and on the right, the gene expression levels over our investigated samples. Tissue gene expression was classified into being not expressed (90th percentile TPM ≤ 1) in any, 1-3, or more than 3 tissues other than testis, to assess tumor specificity. The number of HLAIp identified per gene is depicted, as well as gene (GENCODE) and sample type. **b** The same as above is plotted also for non-coding source genes identified in lung tissue samples and **c** specifically also for the tumor-specific noncHLAIp identified in lung cancer patient C3N-02289.

Using an in-house curated inventory of human transposable element (TE)-derived protein sequences (from three-frame translations) as reference, we found 88 unique TE-HLAIp in our whole dataset. Some were derived from autonomous TEs, such as long tandem repeat (LTR) retrotransposons and long interspersed nuclear elements (LINEs), and others from non-autonomous retrotransposons such as short interspersed nuclear elements (SINE) or SINE-VNTR-Alu (SVA) elements (**Supplementary Fig. 5a**). Importantly, 60 of the 88 TE-HLAIp were found in presumed non-coding TE regions and therefore represent completely novel HLA peptides. These TE-HLAIp would have been overlooked in canonical MS-based searches. Furthermore, 10% of the noncHLAIp derived from TEs were retrospectively shown to be expressed only in a single healthy tissue, excluding testis. For example, peptides derived from AluSq2 SINE/Alu and L1PA16 LINE/L1 elements were expressed only in skin and testis. Finally, selected TAAs were also investigated in the same manner for melanoma and lung tissue samples separately (**Supplementary Fig. 5b-c**).

We next examined whether our approach could identify tumor specific non-canonical targets in the ideal case where normal and tumor biopsies are available, i.e. from the two lung cancer patients included in the present dataset. For example, out of 14,120 non-canonical genes expressed in patient C3N-02289, we found that 409 were exclusively expressed at the RNA level in the patient’s tumor. We identified 45 noncHLAIp in patient C3N-02289 (Fig. 5b), out of which 10 peptides were identified only in the tumor tissue by MS (Fig. 5c). Four of these source genes were tumor-specific when compared to the matched healthy tissue of patient C3N-02289 (90^th^ percentile Transcripts per Million (TPM) ≤ 1). However, when compared to the GTEx database, only one noncHLAIp from RP11-566H8.3 was finally considered as tumor and testis-specific for patient C3N-02289 (Fig. 5c). The same analyses with TE genes resulted in the identification of 1,159 elements that were expressed at the RNA level only in the C3N-02289 tumor. Of those, we identified the LTR7B LTR/ERV1 TE HLAIp that was presented in the tumor tissue, however this gene is also expressed in healthy brain. In comparison, we were able to identify six tumor-associated protHLAIp, which were only found in the tumor tissue (BIRC5, TERT, FAP, SPAG4, MAGEA9 and BCL2L1).

### NoncHLAIp are shared across patient samples

We investigated the likelihood of shared noncHLAIp amongst the nine tumor samples analyzed and identified 27 peptides that were detected in at least two patient samples. Seven noncHLAIp, already validated in 0D5P, were confirmed by PRM in at least one other patient sample that expressed HLA allotypes with identical or highly similar binding specificities (**Supplementary Table 4**), with a total of 15 individually detected PRM events (Fig. 6a). Interestingly, one noncHLAIp VTDQASHIY, derived from microcephalin-1 antisense RNA (MCPH1-AS1), was confirmed independently with PRM in three melanoma or lung cancer patients (**Supplementary Fig. 6a-b**). Further, the shared presentation of noncHLAIp AAFDRAVHF, derived from the family of LINEs (LINE/L2) on chromosome 6, was confirmed in two melanoma samples (**Supplementary Fig. 6c-d**). Interestingly, the corresponding source RNA expression is skin- and testis-restricted.

**Fig. 6.**
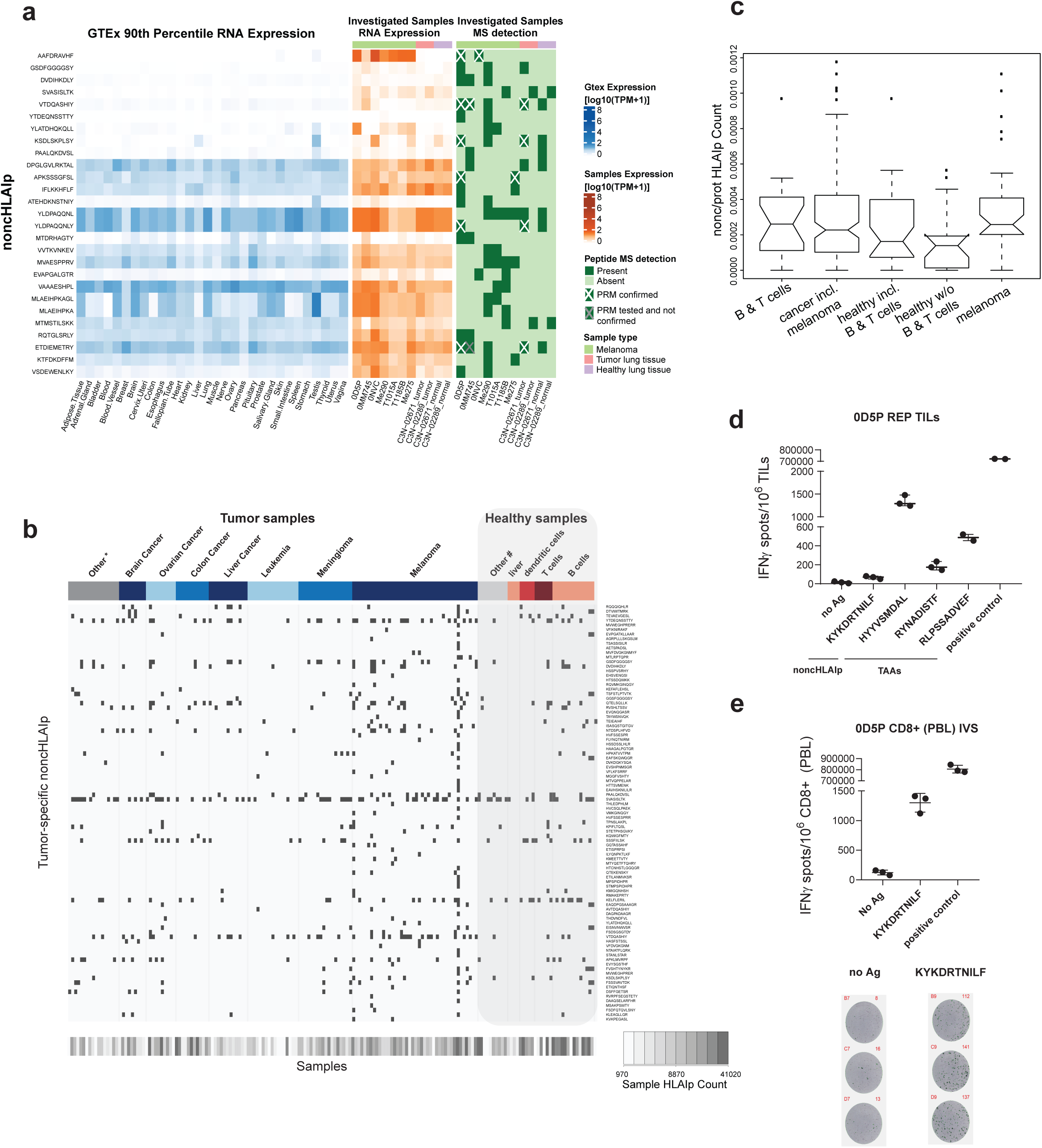
NoncHLAIp can be shared across individuals and are immunogenic. **a** The noncHLAIp-centric heatmap (left) shows the corresponding **non-coding** gene expression (90th percentile) across healthy tissues, as well as in our investigated samples (middle). The peptides that were MS-identified across the investigated samples, and therefore shared, are outlined in the rightmost heatmap. Validation by PRM was performed for multiple noncHLAIp across the corresponding samples and denoted with cross markings. **b** noncHLAIp identification across a large collection of immunopeptidomics datasets (ipMSDB) consisting of both tumor and healthy samples. Tumor specific noncHLAIp were re-identified and were significantly enriched in tumor samples. Tumor samples are labelled in shades of blue, * include tumor metastasis, myeloma, prostate, uterine, kidney, lung and pancreatic cancer, and neuroblastoma. Healthy samples are indicated in shades of red, # include fibroblast cells, monocytes, pancreatic tissue, epithelial cells, normal lung tissue, and apheresis samples. B and T cells mostly constitute of samples that were immortalized or rapidly expanded in culture. **c** Boxplot depicting the number of noncHLAIp identified in the different groups of samples derived from ipMSDB. **d** Reactivity was measured in melanoma 0D5P by IFNγ ELISpot using autologous REP TILs. Three TAAs from TYR and TYRP1, and one non-canonical dORF derived HLAIp from ABCB5, induced an IFNγ response. **e** In addition, CD8+ T lymphocytes from PBLs were re-challenged with autologous CD4+ blasts together with 1μM of the non-canonical ABCB5 HLAIp. (No Ag: no peptide, positive control: 1x cell stimulation cocktail)

Next, we interrogated a large collection of immunopeptidomic datasets (ipMSDB^48^, 137 biological cancer tissue/cell line sources, 39 biological healthy tissues/cell line sources; 2,250 MS raw files in total) and obtained the first large-scale signature of noncHLAIp presentation. In total, 398,622 peptides were obtained for the healthy samples in ipMSDB, versus 488,500 peptides for cancer samples. We observed that 92 out of the 96 tumor-specific noncHLAIp (90^th^ percentile TPM ≤ 1 in maximum 3 tissues) identified in this present study were re-identified in ipMSDB (Fig. 6b), 52 of those were detected in at least one additional sample in ipMSDB excluding our investigated samples. Another 72 additional novel noncHLAIp were discovered from the same tumor specific non-canonical source genes. Remarkably, we observed that noncHLAIp presentation was significantly enriched in tumor samples in ipMSDB (cancer versus healthy p-value=0.048, melanoma versus healthy p-value=0.025), and more significantly when B and T cells, which were rapidly expanded in culture or EBV-transformed, were excluded from the analysis (p-value = 0.009) (Fig. 6c). Interestingly, two noncHLAIp from HAGLROS (KVLAGTVLFK and VLAGTVLFK), identified in the lung cancer tissue only, were exclusively found only in cancer samples in ipMSDB, mainly in ovarian cancer samples, where genetic association with HAGLROS was previously reported^49^.

### Assessing immunogenicity of noncHLAIp with autologous T cells

The involvement of the noncHLAIp in tumor immune recognition was assessed by measuring IFNγ release by autologous tumor-infiltrating lymphocytes (TILs) or peripheral blood mononuclear cells (PBMCs) upon peptide stimulation. Out of the 786 peptides screened (94 TEs, 421 lncRNAs, 56 alternative ORFs, 215 TAAs), we confirmed the specific recognition of autologous TILs to TAAs HYYVSMDAL and RLPSSADVEF from tyrosinase (TYR), and RYNADISTF from tyrosinase-related protein 1 (TYRP1) in melanoma sample 0D5P, and TAA YLEPGPVTA from PMEL in melanoma sample T1015A. One non-canonical peptide KYKDRTNILF, derived from the downstream ORF (dORF) of the melanoma stem cell marker ABCB5 gene in 0D5P, was found to be immunogenic in both CD8+ TILs and CD8+ T cells from peripheral blood lymphocytes (PBLs) (Fig. 6d-e). Notably, this peptide is shared across three additional melanoma samples in ipMSDB.

## Discussion

Our proteogenomics approach led to the stringent identification of hundreds of noncHLAIp derived from non-coding genes, TEs and alternative ORFs. This was achieved with NewAnce, a novel computational module which overcame the challenge of reduced sensitivity and specificity when searching against large MS search spaces^29, 50^. NewAnce is publically available and can be used with any (non-canonical) protein sequence databases of interest. We rigorously tested the validity of noncHLAIp identifications with HLA binding predictions, sequence-specific retention characteristics, targeted MS analyses, and provided evidence of translation in peptide-encoding ORFs by Ribo-Seq. We confirmed with these multiple strategies that NewAnce is superior to MaxQuant and Comet alone, across all investigated samples. As an example with one patient, we conducted PRM and Ribo-Seq analyses to compare a subset of protHLAIp to non-canonical antigen classes (lncRNAs and TEs), thereby validating at the experimental level a recurrently identified noncHLAIp. We found that noncHLAIp had an overall lower confirmation rate than protHLAIp, possibly due to their lower expression, which also led to their stochastic detection by MS. Interestingly, the expression and translation of microproteins derived from presumed non-coding RNAs in the heart were recently discovered using a Ribo-Seq directed proteogenomics approach, with evidence of translation in the correct ORF confirmed for 22.5% of the lncRNAs and 55.4% of the micropeptides validated by PRM MS^51^. Importantly, our results additionally demonstrate that the correct identification of noncHLAIp in proteogenomic workflows requires proper FDR controls and validation using multiple independent methods.

Combining immunopeptidomics with RNA-Seq and Ribo-Seq datasets enables the comprehensive assessment of how transcription, translation and HLA presentation are correlated. Despite the different methodological challenges^52–54^, the expected correlations were observed between HLA presentation level and expression based on both RNA-Seq and, more clearly on Ribo-Seq, perhaps because translation is biologically closer to antigen processing and presentation. In addition, we found that in melanoma 0D5P, most of the novel noncHLAIp derived from the Ribo-Seq inferred database originated from source genes harboring upstream ORFs (uORFs). Of note, uORFs can trigger non-sense-mediated decay of mRNAs and provide a rich source of noncHLAIp^55–57^.

While a previous study has shown that the presentation of non-canonical peptides is enhanced by inflammatory stimuli, only the presentation of specific HLA peptides were documented^58^. In contrast, our large-scale analyses of both decitabine and IFNγ-treated cells did not detect profound changes in noncHLAIp presentation, although non-canonical source genes were induced. Hence, we hypothesize that low copy number of such noncHLAIp still remains a limiting factor for their presentation. Moreover, corroborating prior research^28^, we report the enrichment of noncHLAIp originating from the C-termini of source protein sequences. Translation products of such presumed non-coding regions could be considered as defective ribosomal products that are expected to be unstable and rapidly degraded, likely bypassing the proteasome^59^.

Given the lack of comprehensive healthy tissue immunopeptidomics libraries from patients, we propose a workflow to retrospectively filter for tumor-specific noncHLAIp with publicly available RNA-Seq databases (such as GTEx^46^). We further validated our proteogenomics approach in the ideal situation where both tumor and matched normal tissue are available. Two overlapping epitopes were identified in lncRNA HAGLROs, which were expressed and presented only in the lung tumor tissue. This lncRNA has been implicated in cancer progression^60, 61^ and should now be prioritized for downstream validation. Moreover, while Laumont et al.^24^ first proposed the existence of shared noncHLAIp, our work validates that noncHLAIp can be shared across multiple tumor samples and thus we anticipate greater efficiency in treatment with such shared noncHLAIp compared to private neoantigens^62, 63^.

Expression of tumor-specific noncHLAp in a subpopulation of tumor cells suggests a dependency on a molecular or functional state. For example, the immunogenic noncHLAIp, derived from dORF in the ABCB5 gene was moderately expressed in only 37% of the melanoma cells, compared to the immunogenic TYR and TYRP1, both of which were highly and uniformly expressed. Although targets for immunotherapy should ideally be uniformly presented in all cancer cells in order to minimize outgrowth of escaping cells, immune pressure on selected tumor cell subsets of particular biological relevance – such as cancer stem-like cells, tumor cells with epithelial-mesenchymal transition features or proliferating tumor cells – could affect tumor behavior and be clinically beneficial. Ideally, such targets would also be upregulated by inflammatory cytokines or pharmacologically.

Indeed, we found such immunogenic noncHLAIp from 0D5P derived from the dORF of the ABCB5 gene. ABCB5 has been shown to be expressed in malignant-melanoma-initiating cells and was suggested to be responsible both for the progression and chemotherapeutic refractoriness of advanced malignant melanoma^43^. Through an IL1β/IL8/CXCR1 cytokine signaling circuit, it has been shown to control IL1β secretion and maintain slow cycling and chemoresistance^64^. Blockage of ABCB5 reversed resistance to multiple chemotherapeutic agents, induced cellular differentiation, and impaired tumor growth *in vivo*^64^. We found that ABCB5 was differentially co-expressed in a cluster of 0D5P cells with the transcription factor MITF and beta-catenin, and others that their expression may be enriched in melanoma stem cell populations^65^. The presence of spontaneous specific T cells recognizing the noncHLAIp derived from the dORF of the ABCB5 gene, in both peripheral blood and TILs, suggests no central tolerance, and that this target could allow immune-targeting of melanoma stem cell subpopulation to drastically affect tumor growth.

Out of 571 noncHLAIp screened, immune recognition by rapidly expanded TILs and PBMCs was detected for only a single immunogenic noncHLAIp. Various mechanisms could account for this lack of recognition. First, we were only able to screen autologous TILs that had been long propagated in culture. We have previously reported that TIL *ex vivo* expansion may lead to depletion of T cell clones that recognize tumor neoantigens^66^. Second, it is possible that the melanoma cells, which had to be expanded considerably in culture for immunopeptidomics analyses, could have altered their HLA peptide repertoire, leading to the identification of noncHLAIp that were originally not present in freshly extracted cells. However, we also interrogated snap-frozen lung cancer tissues and still could not detect immune recognition of identified non-canonical targets in autologous PBMCs. Alternatively, the ability of noncHLAIp to induce a natural immune response might be inferior to protHLAp. Low expression might limit uptake by professional antigen presenting cells and hence priming of naïve T cells *in vivo* through cross presentation. Similarly, engagement of CD4+ T helper cells through HLA class II presentation might be limited as well. Nevertheless, tumor-specific non-canonical targets may still be valuable for cancer vaccines, even when no prior immune response against the targets has been detected *ex vivo*, as previously shown for neoantigens^4, 67, 68^. More research should be performed to thoroughly assess the ability of noncHLAp to augment protective immune response *in vivo*. Such approaches are supported by evidence in mouse models that peptides derived from non-canonical regions can be spontaneously recognized and leveraged in cancer immunotherapy^24, 69^.

Remarkably, across tumor types, the potential number of predicted noncHLAp is orders of magnitude larger than that of neoantigens encompassing non-synonymous somatic mutations. As T cell-based screenings currently have limited throughput and are expensive^70^, an accurate and cost-effective non-canonical target discovery pipeline is crucial for their further development and use in cancer immunotherapy. With the renewed interest in cancer vaccines and the constantly growing number of antigens screened for immune recognition, we expect that enough training data will become available to allow the development of accurate predictors of immunogenicity. Combining this with our newly developed module NewAnce to shortlist *in vivo* presented noncHLAp and to rank them according to their predicted immunogenicity, will facilitate the comprehensive exploration of non-canonical antigens, their association with immune responses and their potential for building effective cancer immunotherapies.

## Methods

### Patient material

Melanoma cell lines (0D5P, 0MM745, 0NVC) were generated as follows: Patient-derived tumors were cut into small pieces before being transferred into a digestion buffer containing collagenase type I (Sigma Aldrich) and DNase I (Roche) for at least one hour. Dissociated cells were washed and maintained in RPMI 1640 + GlutaMAX medium (Life Technologies) supplemented with 10% heat-inactivated FBS (Dominique Dutscher) and 1% Penicillin/Streptomycin Solution (BioConcept). If fibroblasts appeared, they were selectively eliminated with G418 (Geneticin; Gibco) treatment. The primary melanoma cell lines T1185B, T1015A, Me290 and Me275 were generated at the Ludwig Cancer Research Center, Department of Oncology, University of Lausanne, as previously described^71, 72^. All established melanoma cells were subsequently grown to 1 x 10^8^ cells, collected by centrifugation at 151 x g for 5 min, washed twice with ice cold PBS and stored as dry cell pellets at −20°C until use. For the i*n vitro* 72h treatment with IFNγ (100 IU/mL, Miltenyl Biotec), T1185B cells were grown to 2 x 10^8^ in triplicates. For the treatment with Decitabine (DAC, Sigma-Aldrich), 2 x 10^8^ melanoma cells were grown for 8 days in 0.5 µM Decitabine with medium and drug renewal on the 4th day.

Autologous TILs were expanded from fresh melanoma tumor samples corresponding to patients 0D5P, 0MM745, 0NVC, LAU1185 (tumor cell line T1185B), LAU1015 (tumor cell line T1015A), LAU203 (tumor cell line Me290) and LAU50 (tumor cell line Me275) at the Ludwig Cancer Research Center, Department of Oncology, University of Lausanne. The fresh tissues were manually cut into fragments of one to two mm^3^. The tumour fragments were then placed in 24-well plates containing RPMI CTS grade (Life Technologies), 10% Human serum (Biowest), 0.025 M HEPES (Life Technologies), 55 μmol/L 2-Mercaptoethanol (Life Technologies) and supplemented with IL-2 (6,000 IU/mL, Proleukin) for three to five weeks. Following this pre-REP (Rapid Expansion Protocol), TILs were then expanded with another REP as follows: 5×10^6^ TILs were stimulated with irradiated feeder cells (Ratio 1:200), anti-CD3 (OKT3, 30 ng/mL, Miltenyl biotec) and IL-2 (3,000 IU/mL) for 14 days. After 14 days of REP, about 2×10^9^ TILs were harvested, washed and cryopreserved until use. The purity (i.e. the % of CD3 T cells) was >95%. As additional control, one flask with the exact same REP conditions without TILs was cultured in parallel and no cells were detectable at day 14. REP TILs were thawed in 5 IU/mL DNAse I (Sigma-Aldrich) and cultured in 3000 IU/mL IL-2 for two days in RPMI 1640 Medium with GlutaMAX™ Supplement (Gibco), with the addition of 8% Human serum (Biowest), 10 mM HEPES (Gibco), 50 μM Beta-Mercaptoethanol (Gibco), 100 μM non-essential amino acids (Gibco), 100 IU/mL Penicillin and 0.1 mg/mL Streptomycin, 2mM L-Glutamine (Gibco), 0.1 mg/mL Kanamycinsulfate (Carl Roth) and 1mM sodium pyruvate (Gibco). Cells were then washed twice in complete medium and subsequently rested overnight in the presence of 150 IU/mL IL-2 prior to peptide stimulation.

Snap-frozen normal and lung tumor tissue material from C3N-02289 (Lung squamous cell carcinoma, grade 2) and C3N-02671 (Lung adenocarcinoma, G2) were kindly provided by the International Institute of Molecular Oncology. Informed consent of the participants was obtained following requirements of the institutional review board (Ethics Commission, CHUV, Bioethics Committee, Poznan University of Medical Sciences, Poznań, Poland).

All cells were tested negative for mycoplasma contamination. High resolution 4-digit HLA-I and HLA-II typing (**Supplementary Table 1**) was performed either at the Laboratory of Diagnostics, Service of Immunology and Allergy, CHUV, Lausanne or in-house with the following method: The amplification of the HLA was conducted with the TruSight HLA v2 Sequencing Panel kit (CareDx) according to the manufacturer’s protocol. Sequencing was performed on the Illumina® MiniSeq™ System (Illumina) using paired-end 2×150 bp protocol. The data was analyzed with the Assign TruSight HLA v2.1 software (CareDx).

### Immunoaffinity purification of HLA peptides

We performed HLA immunoaffinity purification following our previously established protocols^39, 73^. Briefly, W6/32 and HB145 monoclonal antibodies were purified from the supernatant of HB95 (ATCC® HB-95™) and HB145 cells (ATCC® HB-145™) using protein-A sepharose 4B (Pro-A) beads (Invitrogen), and antibodies were then cross-linked to Pro-A beads. Cell lysis was performed with PBS containing 0.25% sodium deoxycholate (Sigma-Aldrich), 0.2 mM iodoacetamide (Sigma-Aldrich), 1 mM EDTA, 1:200 Protease Inhibitors Cocktail (Sigma-Aldrich), 1 mM Phenylmethylsulfonylfluoride (Roche), 1% octyl-beta-D glucopyranoside (Sigma-Alrich) at 4°C for 1 hour. Lysates were cleared by centrifugation in a table-top centrifuge (Eppendorf) at 4°C at 21,191 x g for 50 min. Snap-frozen tissue samples were homogenized on ice in 3-5 short intervals of 5 seconds each using an Ultra Turrax homogenizer (IKA) at maximum speed. Lysates were cleared by centrifugation at 25,000 rpm in a high speed centrifuge (Beckman Coulter, JSS15314) at 4°C for 50 minutes. We employed the Waters Positive Pressure-96 Processor (Waters) and 96-well single-use micro-plates with 3µm glass fiber and 10µm polypropylene membranes (Seahorse Bioscience, ref no: 360063). Anti-pan HLA-I and HLA-II antibodies cross-linked to beads were loaded on their respective plates. For tissue samples, a depletion step of endogenous antibodies was required containing Pro-A beads. The lysates were passed sequentially through HLA-I and -II plates at 4°C. Plates were then washed separately with varying concentrations of salts using the processor. Finally, beads were washed twice with 2 mL of 20 mM Tris-HCl pH 8.

Sep-Pak tC18 100 mg Sorbent 96-well plates (Waters, ref no: 186002321) were used for the purification and concentration of HLA-I and HLA-II peptides. Plates were conditioned and direct elution of the HLA complexes and the bound peptides from the affinity plate with 1% trifluoroacetic acid (TFA; Sigma-Aldrich) was performed. After washing the C18 wells with 2 mL of 0.1 % TFA, HLA-I peptides were eluted with 28% Acetonitrile (ACN; Sigma-Aldrich) in 0.1% TFA. HLA-II peptides were eluted from the class II C18 plate with 500 µL of 32% ACN in 0.1% TFA. Recovered HLA-I and -II peptides were dried using vacuum centrifugation (Concentrator plus, Eppendorf) and stored at −20°C.

### LC-MS/MS analyses

The LC-MS/MS system consists of an Easy-nLC 1200 (Thermo Fisher Scientific) hyphenated with a Q Exactive HF-X mass spectrometer (Thermo Fisher Scientific). Peptides were separated on a 450 mm analytical column of 75 µm inner diameter.

For HLAIp, we used the following gradient for analytical separation, with a flow rate of 250 nL/min using a mix of 0.1 % FA (solvent A) and 0.1% FA in 80 % ACN (solvent B): 0–5 min (5% B); 5-85 min (5-35% B); 85-100 min (35-60 % B); 100-105 min (60-95% B); 105-110 min (95% B); 110-115 min (95-2% B) and 115-125 min (2% B). For HLAIIp, the gradient was run as follows: 0-5 min (2-5% B); 5-65 min (5-30% B); 65-70 min (30-60% B); 70-75 min (60-95 % B); 75-80 min (95% B), 80-85 min (95-2% B) and 85-90 min (2% B).

The mass spectrometer was operated as follows for discovery data-dependent acquisition (DDA). Full MS spectra were acquired in the Orbitrap from m/z = 300-1650 with a resolution of 60’000 (m/z = 200) and ion accumulation time of 80 ms. The auto gain control (AGC) was set to 3e6 ions. MS/MS spectra were acquired in a data-dependent manner on 10 most abundant precursor ions (if present) with a resolution of 15,000 (m/z = 200), ion accumulation time of 120 ms and an isolation window of 1.2 m/z. The AGC was set to 2e5 ions, dynamic exclusion to 20 s and a normalized collision energy (NCE) of 27 was used for fragmentation. No fragmentation was performed for HLAIp in case of assigned precursor ion charge states of four and above, and for HLAIIp, in case of assigned precursor ion charge states of one, and also from six and above. The peptide match option was disabled.

### Parallel Reaction Monitoring

Selected endogenous HLAp that required confirmation by parallel reaction monitoring (PRM) were ordered as crude (PePotec grade 3) or HPLC grade (purity > 70%) with one stable isotope labelled amino acid from Thermo Fisher Scientific. The mass spectrometer was operated at a resolution of 120’000 (at m/z = 200) for MS1 full scan, scanning a mass range from 300-1650 m/z with an ion injection time of 100ms and an AGC of 3e6. Then each peptide was isolated with an isolation window of 2.0 m/z prior to ion activation by HCD (NCE = 27). Targeted MS/MS spectra were acquired at a resolution of 30’000 (at m/z = 200) with an ion injection time of 60ms and an AGC of 5e5. Only those peptides that ultimately pass quality control were considered for further downstream analyses through spiking them back into the patient sample.

The PRM data was processed and analyzed by Skyline (v4.1.0.18169, MacCoss Lab Software)^74^ and an ion mass tolerance of 0.02 m/z was used to extract fragment ion chromatograms. To display MS/MS spectra, raw data was converted into MGF by MSConvert (Proteowizard v3.0.18136) and peak lists for the heavy peptide and light counterpart were extracted. The assessment of MS/MS matching was performed by pLabel (Version 2.4.0.8, pFind studio, Sci. Ac.) and Skyline.

### Exome/RNA sequencing

DNA was extracted for HLA typing and exome sequencing with the commercially available DNeasy Blood & Tissue Kit (Qiagen), following manufacturers’ protocols. For tissue samples, pelleted DNA was used, which was obtained after lysis of the tissue and centrifugation during HLA immunopurification. The supernatant was used for HLA immunopurification, whereas the pelleted DNA was homogenized with a pestle (70 mm, 1.5/2.0mL, Schuett-Biotec) before DNA extraction following manufacturer’s instructions.

RNA extraction for RNA sequencing was achieved using the total RNA isolation RNeasy Mini Kit (Qiagen) following manufacturer’s protocols for all melanoma cell lines (including DNAse I (Qiagen) on-column digestion). Frozen pieces of tumor and normal tissue samples (<20 mg) were directly submerged in 350 µL of RLT buffer supplemented with 40 µM DTT (Sigma-Aldrich). Tissues were then completely homogenized on ice using a pestle (70 mm, 1.5/2.0mL, Schuett-Biotec) and by passing the sample through a 26G needle syringe for five times (BD Microlance). Centrifugation was performed in a table-top centrifuge (Eppendorf) at 4°C at 18,213 x g for 3 min, before the supernatant was removed and used for RNA extraction. All subsequent steps are described in detail in the manufacturer’s protocol (including DNAse I (Qiagen) on-column digestion).

Three micrograms of genomic DNA were fragmented to 200bp using a Covaris S2 (Covaris). Sequencing libraries were prepared with the Agilent SureSelectXT Reagent Kit (Agilent Technologies). Exome enrichment was performed with Agilent SureSelect XT Human All Exome v5 probes. Cluster generation was performed with the resulting libraries using the Illumina HiSeq PE Cluster Kit v4 reagents and sequenced on the Illumina HiSeq 2500 using SBS Kit v4 reagents. At least 70x coverage for the melanoma cell lines and PBMCs/TILs were required. For tumor/normal lung tissues, at least 100x coverage was required. Sequencing data were demultiplexed using the bcl2fastq Conversion Software (v. 1.84, Illumina).

RNA quality was assessed on a Fragment Analyzer (Agilent Technologies) and all RNA had a RQN beween 7.4 and 10. RNA-seq libraries were prepared using 500 ng or 375 ng of total RNA with the Illumina TruSeq Stranded mRNA reagents (Illumina) following manufacturer’s recommendations. Libraries were quantified by a fluorimetric method and their quality assessed on a Fragment Analyzer (Agilent Technologies). Cluster generation was performed from the resulting libraries using the Illumina HiSeq PE Cluster Kit v4 reagents and sequenced on the Illumina HiSeq 2500 using HiSeq SBS Kit v4 paired end reagents for 2×100 cycles paired end sequencing. Sequencing data were de-multiplexed using the bcl2fastq2w Conversion Software (v. 2.20, Illumina).

### RNA-Seq data processing for lncRNA and gene expression analysis

The GENCODE comprehensive gene annotation version 221,2 was downloaded from the GENCODE website (https://www.gencodegenes.org/releases/22.html). It was used to define the protein-coding and non-coding gene features including chromosome position, transcript structure, as well as transcript and protein sequences. Here, the human reference genome GRCh38/hg38 was downloaded from the UCSC Genome Browser website (http://hgdownload.cse.ucsc.edu/goldenPath/hg38/bigZips/) and was used as the genome assembly. The RNA-seq reads were aligned to the GRCh38/hg38 reference genome using RNA-Star (v2.4.2a; https://github.com/alexdobin/STAR). The gene expression was normalized and calculated as in Fragments Per Kilobase of transcript per Million mapped reads (FPKM) values by Cufflinks (v2.2.1) (http://cole-trapnell-lab.github.io/cufflinks/releases/v2.2.1/). The gene level RNA expression data of both protein-coding and non-coding genes were used for downstream gene expression analysis^33, 75^.

### RNA-Seq data processing for TE expression analysis

Reads from the investigated samples and public data from GTEx were mapped to the human (GRCh37) genome using hisat2 v.2.1.0^76^. Counts on genes and TEs were generated using featureCounts 1.6.2^77^. To avoid read assignation ambiguity between genes and TEs, a gtf file containing both was provided to featureCounts. For repetitive sequences, an in-house curated version of the Repbase database was used (fragmented LTR and internal segments belonging to a single integrant were merged). Only uniquely mapped reads were used for counting on genes and TEs. Finally, features that did not have at least one sample with 20 reads were discarded from the analysis. Normalization for sequencing depth was done for both genes and TEs using the TMM method as implemented in the limma v.3.36.5 package of Bioconductor^78^ and using the counts on genes as library size.

### Personalised sequence databases for non-coding genes

The curated set of human ENCODE non-coding transcripts (GRCh37 reference assembly) was downloaded from https://www.gencodegenes.org/human/release_24lift37.html. Short ORFs (shORFs) in all three forward reading frames were identified using a stop-to-stop strategy. The minimum peptide length was set at 8 amino acids, and the longest polypeptide identified was 3644 amino acids. Unless otherwise mentioned, all non-coding genes that were expressed per sample (FPKM > 0) were translated in all three forward reading to build the personalized fasta file for non-coding genes.

### Personalised sequence databases for protein-coding genes including variants

GENCODE v24 (GRCh37 human reference assembly, downloaded from https://www.gencodegenes.org/human/release_24lift37.html) was chosen as the standard reference dataset. Whole exome sequence reads were aligned to the GRCh37 human assembly with BWA-MEM^79^ and variants were predicted using the GATK framework v3.7 and Picard Tools v 2.9.0^80^. SNPs were defined as variants present in both tumor and germline, and somatic mutations (SNVs and indels) as present only in tumor. The GENCODE comprehensive gene annotation file, in GFF3 format, was parsed to extract genomic coordinate information for every exon in each protein coding transcript, and those coordinates were compared with sample-specific variant coordinates to derive non-synonymous amino acid changes within each protein. For every sample, we created a separate fasta file where residue mutation information was added to the header of the affected translated protein coding transcripts, in a format compatible with MaxQuant v1.5.9.4i as previously reported^81^.

### Mass Spectrometry Database Search

We used two widely used search tools: Comet 2017.01 rev. 2^35^ and the Andromeda search engine within MaxQuant v1.5.9.4i^82^. Both Andromeda and Comet allow searching for peptides with and without variants. Andromeda matched the MS/MS spectra of each sample against their personalized reference libraries (mentioned above). Similarly, the variants were annotated in the PEFF format (http://www.psidev.info/peff) for Comet. Both search tools were run with the same principal search parameters: precursor mass tolerance 20ppm, MS/MS fragment tolerance 0.02 Da, peptide length 8-15 for HLA-I only and 8-25 for HLA-I and HLA-II peptides and no fixed modifications. For samples 0D5P, 0VNC and OMM745, oxidation (M) and phosphorylation (STY) were set as variable modifications and for the remaining samples only Oxidation (M) was included as a variable modification. A PSM FDR of 3% was used for Andromeda as a first filter, and non-canonical reference sequences were loaded into the “proteogenomics fasta files” module for separate FDR calculations for protein-coding and non-canonical sequences.

To assure that non-canonical peptide sequences do not match other protein coding genes, all peptides found by Andromeda or Comet were aligned against the UniProt (www.uniprot.org) sequence database (human reviewed sequences with isoforms, downloaded 18/12/2018), where leucine and iso-leucines were treated as equal since they are not distinguishable by mass spectrometry. If peptides were found matching standard UniProt sequences, they were assigned as protein-coding with the UniProt IDs. However, we assigned and kept TE peptide sequences that matched annotated TEs that were integrated into the human reference in UniProt.

As schematically described in **Supplementary Fig. 1a**, the Comet FDR calculation was done separately for protein-coding and non-canonical PSMs with an in-house software written in Java, which utilizes the MzJava class library ^83^. All PSMs resulting from the Comet search, including the decoy PSMs (decoy hits originating from reversed sequences), were split into three sublists with PSMs of charge (Z) 1, 2, and charge 3 or higher. The three Comet scores XCorr, deltaCn and spScore were considered. It has been shown that when feature vectors are partitioned into different groups, group-wise local FDR (lFDR) calculation provides the most sensitive decision boundaries, for controlling the global FDR^84^. Therefore, the 3D space (XCorr, deltaCn and spScore) was partitioned into small cells (40 intervals in each dimension) and the lFDR was estimated for each cell. We used the following equation to calculate the lFDR:

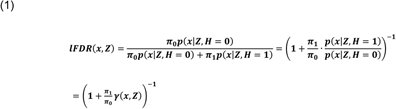

where ***π*_0_** and ***π*_1_** are the class probabilities for true (H=1) and wrong (H=0) PSMs, and ***p*(*x*|*Z*, *H* = 0, 1)** are the probability distributions for feature vector *x* = (***XCorr, deltaCn, spScore***) of PSMs with charge ***Z***. The probability ratio ***γ*(*x*, *Z*)** is estimated for every cell and charge using all the target and decoy PSMs. The **π_1_⁄π_0_** ratios are calculated for the non-canonical and protein-coding groups separately and then the **π_1_⁄π_0_** ratios are plugged into Equation (1). This way the lFDR values are calculated for every cell for both groups. For each cell *i*, the number of wrong hits (*n*_0i_) is set to the number of decoy hits, while the number of true hits (*n*_1i_) is set to the number of target hits minus *n*_0i_. These counts are then smoothed by averaging over the neighboring cells. Then ***γ(x,Z)* = *n_1i_ · n_0,tot_/n_0i_·n_1,tot_*** or ***γ*(*x*, *Z*) = 1** if *n*_0i_ = 0, where *n*_0,tot_ and *n*_1,tot_ are the total number of decoy and target hits in all cells. The target and decoy PSMs are further split into non-canonical and protein-coding PSMs. Since there usually are not enough PSMs to estimate ***γ*(*x*, *Z*)** for the non-canonical group, ***γ*(*x*, *Z*)** is taken from all PSMs but **π_1_⁄π_0_** is adapted for each group. Therefore, it is assumed that the score distributions are the same for each group, but that the ratio of true to wrong PSMs may change. The **π_1_⁄π_0_** ratios are calculated for the non-canonical and protein-coding groups separately and then the **π_1_⁄π_0_** ratios are plugged into Equation (1). This way the lFDR values are calculated for every cell for both groups separately. The **π_1_⁄π_0_** ratio in the non-canonical group is relatively smaller than the protein-coding **π_1_⁄π_0_** ratio, because the non-canonical database is larger and mostly consists of peptides that are not present in the sample. This will lead to a larger non-canonical lFDR value, and to a more conservative filtering of non-canonical PSMs. Finally, the lFDR threshold is adjusted to allow a global FDR of 3% for the non-canonical and protein-coding groups.

PSMs from both search tools were combined and only the intersection, meaning PSMs with identical Comet and Andromeda matches (same peptide sequence with the same modification) were retained. In order to assign peptides into source protein groups, we implemented a greedy bipartite graph protein grouping algorithm^85^. The total and ‘unique’ peptide counts were calculated for each protein. To calculate the adjusted peptide counts we sorted the proteins in each group by decreasing number of peptides and for each protein removed the peptides of all proteins higher up in the list.

To build the ipMSDB database, we searched 2,250 immunopeptidomics raw files with Comet (PSM FDR of 1%, as described above), and the Apache Spark cluster computing framework^86^ was used to process the results and calculate the FDR. The samples were annotated with basic biological information for further statistical analysis.

### Ribo-Seq: Experimental Protocol

Ribo-Seq was performed according to Calviello et al 2016^87^. Ribo-Seq libraries were derived from 80% confluent 10 cm tissue culture dishes of adherent melanoma 0D5P cells. Following a wash with ice-cold PBS supplemented with 100 μg/mL cycloheximide (Sigma Aldrich), the cells were immediately snap-frozen by placing the dishes on liquid nitrogen, which were then placed on wet ice. Lysis Buffer containing 20 mM Tris-Cl pH 7.4, 150 mM NaCl, 5 mM MgCl2, 1 mM DTT (Sigma Aldrich) and 100 μg/mL cycloheximide, 1% (v/v) Triton X-100 (Calbiochem) and 25 U/mL TURBO DNase (Life Tech) at a volume of 400 uL was immediately added to frozen cells. The cells and buffer were then scraped off, mixed by pipetting and transferred to eppendorf tubes and kept to lyse on ice for 10 minutes. The lysate was then titurated through a 26-G needle for 10 times with 1 mL syringes and cleared by centrifugation for 10 min at 20,000 x g at 4°C. The cleared supernatant was then transferred to a pre-cooled tube on ice, and footprinting was performed by adding 1000U of RNase I (Life Tech. #AM2295) per 400 μL of the lysate and incubation in a thermomixer set to 23°C, shaking at 500 rpm for 45 min. The digestion was stopped by adding 13 µL SUPERASE-In (Thermo, 20 U/µL) per 400 µL of lysate.

Ribosomes were recovered using two MicroSpin S-400 HR columns (GE Healthcare) per sample. The columns were first equilibrated with a total of 3 mL of buffer containing 20 mM Tris-Cl pH 7.4, 150 mM NaCl, 5 mM MgCl2 and 1mM DTT by performing 6 rounds of washes with 500 uL of the buffer. The resin was resuspended with the last wash and drained by spinning for 4 min at 600 x g. One half of the sample volume was then filtered per column for 2 min at 600 x g, and the filtered halves were then combined. To the combined flow-through, three volumes of Trizol LS (Life Tech) were added and RNA was extracted using the Direct-zol RNA Mini-Prep kit (Zymo Research) following the manufacturer’s instructions (including DNase I digestion). RNA was finally eluted in 30 μL of nuclease-free water and quantified using the Qubit RNA Broad Range Assay (Life Tech).

Ribosomal RNA depletion was performed from up to 5 μg of footprinted RNA using the RiboZero Magnetic Gold kit (Illumina) following the manufacturer’s protocol. Footprinted RNA was precipitated from the supernatant (90 μL) using 1.5 μL of glycoblue (Life Tech), 9μL of 3 M sodium acetate and 300μL of ethanol by snap-freezing in liquid nitrogen and a one-hour to overnight incubation at −80°C and pelleted for 30 min at 21,000 x g at 4°C. The RNA pellet was dissolved in 10 μL of RNase-free water.

Following rRNA depletion, isolation of short fragments and phosphorylation of these by a T4 PNK treatment, sequencing libraries were prepared using the NEXTflex Small RNA-Seq Kit v3 (BiooScientific). Following the manufacturer’s instructions, adapters were diluted 1:2 to decrease adapter dimerization. To determine the optimal number of PCR cycles for library amplification, a pilot PCR with the respective forward and reverse primers was performed for each sample for 12, 14, 16, 18 and 20 cycles. Adapter and primer sequences are published by BiooScientific. Products were run on a native PAGE and optimal cycle numbers were determined as the threshold cycle of the library product of 160 bp, which is expected size for ribosome protected fragments, showing up on the gel with as little adapter dimer product (130 bp) as possible. After the final PCR, libraries were run on and excised from an agarose gel, followed by clean-up using Zymoclean Gel DNA Recovery (Zymo Research). Library quantification and validation were performed using a Qubit dsDNA HS and Bioanalyzer DNA HS assay, respectively. Three 0D5P control samples and three DAC treated samples (in a pool of 21 libraries) and additionally also two 0D5P samples (in a pool of 3 libraries) were sequenced on a NextSeq 500 machine at a loading concentration of 1.6pM using High Output Kits v2 (Illumina) with 75 cycles single-end.

### Ribo-Seq: Analysis

Ribo-Seq reads were stripped of adaptor sequences using cutdapt, and contaminants such as tRNAs and rRNA were removed by alignment to a contaminants index via Bowtie v 2.3.5, consisting of nucleotide sequences from known human rRNA and tRNA sequences drawn from the GENCODE annotation v24^88^. Unaligned reads from this analysis were then aligned to human genome version hg19 with the STAR v 2.6.1a_08-27^89^ splice-aware alignment tool allowing for up to 1 mismatch. The star genome index was built using GENCODE version 24 (lift 37). Reads with up to 20 multi-mapping positions were included, with multi mapping reads beings separately treated in subsequent periodicity analysis. The RIboseQC pipeline v1.0^90^ was used to deduce P-site positions from Ribo-Seq reads, and this P-site data was then used as input to the SaTAnn pipeline v1.0^91^ in combination with custom R scripts^87^ for ORF-calling. The SaTAnn pipeline searches for the periodic ribosomal footprint pattern characteristic of translated ORFs using a supplied database of transcripts, yielding a set of ORFs corresponding to known coding regions, as well as ORFs originating in untranslated regions, non-coding RNAs, intron retentions, and read-through events. 0D5P samples had a median of 2.8 million reads mapping to coding sequences per sample, which constituted a median of 81% of the total reads. Since the false positive rate of periodicity based ORF calling is expected to be tolerant to non-periodic sources of noise such as genomic contamination, we included all samples for 0D5P. ORFs were called in both individual libraries and in the pooled set of all libraries for 0D5P, and ORFs which were fully contained within ORFs detected in another library were merged. ORFs were tested for periodicity, by a multitaper test^87^ and those with a p-value of below 0.05 were kept for analyses.

Protein sequences in fasta format were generated from the coordinates of these ORFs, and used both for validation of peptides found using the RNA-Seq based database, and as a *de novo* assembled database for the subsequent round of peptide detection. Peptides were considered validated by Ribo-Seq if they matched anywhere within the translated ORF sequences.

Riboseq profile plots were plotted with P-site numbers per-base on a log2 (n+1) scale.

### 10x Genomics pipeline and gene expression analyses

For single-cell library preparation on the 10x Genomics platform, the Chromium Single Cell 3′ Library and SingleCell 3’ Reagent v3 were utilized, following the official user guide CG000183 RevA, and the instrument 10x Chromium single cell controller. A total of 1,692 0D5P cells were captured for single-cell transcriptomics. Resulting cDNA libraries were sequenced on NextSeq v 2.5 (with Illumina protocol #15048776). The Cell Ranger v.3.0.1 software (10x Genomics) (https://support.10xgenomics.com/single-cell-gene-expression/software/pipelines) was used to process data generated using the 10x Chromium platform, with a restriction to include only 1400 cells to avoid cells or debris with low UMI counts. This led to the detection of 19,178 genes with a mean of 125,937 mapped reads. Genes present in at least five cells and cells detecting at least 200 genes but no more than 50% of mito genes were kept for the rest of the analysis. This resulted in a reduced matrix of 15,710 genes over 1,365 cells.

The raw counts were log-normalised using the NormalizeData implemented in the Seurat R package (Seurat v3). Prior to further processing, we scaled the data to remove cell-cell variations due to cell cycling or high percentage of mitochondrial genes. For cell cycling correction, we followed the scoring strategy described in Tirosh et al, 2016^92^: each cell was assigned a “Cell Cycle” score and the difference between G2M and S phase scores was regressed out. The clustering was obtained using a graph-based method implemented in Seurat (FindClusters with a resolution set to 0.5) leading to the identification of 5 clusters. Marker genes for each cluster were identified with FindMarkers from Seurat by setting the logFC threshold parameter to 0.15. Marker genes with an adjusted bonferroni p-value < 0.05 were considered significantly differentially expressed. Functional analyses were performed with STRING-db v11 on each cluster using their corresponding marker genes as input.

### Interrogating T-cell reactivity

Peptides were synthesized and lyophilized by the Protein and Peptide Chemistry Facility at the Ludwig Cancer Research Center (crude - >80% purity), Department of Oncology, University of Lausanne, or by Thermo Scientific, and resuspended in DMSO at 10 mg/mL. IFNγ ELISpot assays were conducted to assess the reactivity of the REP TILs towards antigens of interest (TAAs, noncHLAIp) using pre-coated 96-well ELISpot plates (Mabtech) following the manufacturer’s protocol. If necessary, REP TILs were stimulated *in vitro* for 14 days with a single peptide or peptide pool at 1μg/mL before re-challenging with the peptide to assess IFN γ response. For this purpose, REP TILs were plated at 1-2×10^5^ cells per well and challenged for 18h with cognate peptides at a final peptide concentration of 1 µM, in triplicates. Medium without peptide was used as negative control, and 1x Cell Stimulation Cocktail (eBioscience™, Thermo Fisher Scientific) was used as positive control. Spot-forming units were quantified using the Bioreader-6000-E automated counter (BioSys). Positive hits were identified by having more spots than the negative control wells, which did not contain any peptide, plus 3 times the standard deviation of the negative control. Positivity was confirmed in at least ≥ 2 independent experiments.

Identification of circulating antigen-specific T cells in patient 0D5P was performed as previously described ^66, 93^. CD19+ cells were isolated from cryopreserved PBLs using magnetic beads (Miltenyi) and expanded for 14 days with multimeric-CD40L (Adipogen, Epalinges, Switzerland, 1μg/mL) and IL-4 (Miltenyi, 200 IU/mL). CD8+ T lymphocytes were isolated from cryopreserved PBL using magnetic beads (Miltenyi) and were co-incubated at a 1:1 ratio with irradiated autologous CD40-activated B cells and peptides (single peptide or pools of ≤ 50 peptides, 1 µM each). After 12 days of *in vitro* expansion, CD8+ T cells were re-challenged with cognate peptide and T cell responses were assessed by ELISpot.

### Statistical analyses

All statistical analyses have been indicated where appropriate. The following tools were used for statistical analyses: GraphPad Prism 8, Perseus 1.5.5.3, RStudio 3.5.1 and Python 3.6.

### HLA-binding predictions

In order to evaluate the binding affinity of HLAIp, MixMHCpred.v2 prediction was run on all HLAIp of length 8-14. Peptides with a p-value < 0.05 were considered as binders.

### Sequence Specific Retention Calculator

Sequence specific retention was calculated with an online available tool SSRCalc^94^: http://hs2.proteome.ca/SSRCalc/SSRCalcQ.html. Only unmodified peptides were included. Peptides and their mean retention times were plotted against the predicted hydrophobicity indices, which were obtained from the SSRCalc based on the 100Å C18 column, 0.1% formic acid separation system and without cysteine protection.

### Correlation analyses

Correlative analyses between the immunopeptidome and transcriptome of 0D5P (Fig. 2a**-d)** were achieved by first assigning HLAp to their respective source genes. For noncHLAp, unless otherwise indicated, if the peptide map back to more than one non-coding source gene, the gene with the highest transcript expression was allocated for further analyses.

### Assessing HLAIp sampling

For HLAIp sampling analyses, peptides were assigned to source protein groups as described above. Adjusted peptide counts were taken, summed over a gene, and subsequently matching to their corresponding expression values (either transcriptome or translatome based). Normalized sampling corresponds to the adjusted peptide count per protein, normalized by the protein length. Determination of correlation between gene expression or spectral coefficient of 3-periodic signal in Ribo-Seq data and HLA presentation were assessed by fitting a polynomial curve of degree 3 to each dataset. Pearson correlation was calculated to assess the correlation between the fitted curve and the data.

### Peptide position analysis

For peptide position analysis within a protein sequence (**Supplementary Fig. 2**), protein-coding datasets fitting to the length distribution of the 95% confidence level of the lncRNA dataset were selected. Then, the position of HLAp, relative to the full protein sequence, was calculated for source lncRNA and protein-coding sequences. Since the data was not normally distributed, the Wilcoxon test was chosen for statistical analysis.

### PRM analyses

For analyses of PRM statistics, MS-based intensities were taken from the initial MaxQuant peptide table output. Tumor-associated antigens (TAAs) for PRM and further comparative analyses were selected from a non-exhaustive list of known and clinically relevant TAAs.

### GTEx RNA expression analyses

Tissue-specific gene expression data was downloaded from The Genotype-Tissue Expression (GTEx) project, a public resource that contains data from 53 non-diseased tissues across nearly 1000 individuals^46^. We used a custom R script to retrieve gene expression values, based on GTEx v7 publicly available data. In the case of multiple transcripts matching the same entry, expression data of the most expressed one were used. The 90th percentile expression of the gene in the tissue-derived tumor was reported. Investigated sample’s FPKM expression units were converted into TPM units for the purpose of comparison with GTEx data. The R-package “ComplexHeatmap”^95^ from the Bioconductor suite was used to draw heatmaps.

## Data availability

Sequence data has been deposited at the European Genome-phenome Archive (EGA), which is hosted by the EBI and the CRG, under accession numbers EGAS00001003723 and EGAS00001003724. MS raw files, corresponding fasta reference files and NewAnce outputs have been deposited to the ProteomeXchange Consortium via the PRIDE^96^ partner repository with the dataset identifier PXD013649.

### Acknowledgements

We thank the patients for their participation in this study and the Center of Experimental Therapy CHUV, Lausanne, for the collection, generation and access to the samples. We also thank the Genomic Technologies Facility and Johann Weber from the University of Lausanne for providing the sequencing data. We thank the Gene Expression Core Facility at the *École polytechnique fédérale* of Lausanne for scRNA-Seq data. We would like to also express our thanks to Julien Schmidt and Philippe Guillaume at the Protein and Peptide Facility and Tetramer Core Facility, University of Lausanne for the synthetic peptides. Further, we thank Raphael Genolet and Lise Queiroz for the HLA typing of the samples. Lastly, we are very grateful to Prof. Daniel Speiser and Nicolas Gerstermann for kindly providing the melanoma cell lines Me290, Me275, T1015A and T1185B and autologous REP TILs for this present study.

## Authors’ Contributions

G.C. and M.B.S. conceived and designed the project and interpreted the results. M.B.S. and C.C. designed the experiments, coordinated, integrated, and interpreted the multi-omics analyses. M.M. conceptualized and implemented the software for MS/MS data processing and FDR calculation in NewAnce, and performed ipMSDB analyses. C.C. J.M. and H.S.P. conducted the immunopeptidomics MS experiments. F.H. assisted in data visualization and GTEx analyses. C.C. and A.A. conducted TIL experiments and A.Ha. assisted with data interpretation. B.J.S. performed NGS analyses. I.X provided support for the computational infrastructure. D.G. and E.P. performed bioinformatics quantification and analysis of TE expression and L.S-R. and D.T. provided scientific interpretation of TE analysis. M.L. analyzed scRNA Seq data. L.Z. analyzed RNA seq data for prot and nonc genes. I.B. and A.Hi. processed the samples for Ribo-Seq and D. H., L.C. and U.O. analyzed and interpreted the Ribo-Seq data. M.W., J.L and M.B. coordinated tissue collection and processing from lung cancer patients. J.L. performed pathological evaluation. C.C. and M.B.S. wrote the manuscript with contributions from all authors.

